# CountASAP: A Lightweight, Easy to Use Python Package for Processing ASAPseq Data

**DOI:** 10.1101/2024.05.20.595042

**Authors:** Christopher T. Boughter, Budhaditya Chatterjee, Yuko Ohta, Katrina Gorga, Carly Blair, Elizabeth M. Hill, Zachary Fasana, Adedola Adebamowo, Farah Ammar, Ivan Kosik, Vel Murugan, Wilbur H. Chen, Nevil J. Singh, Martin Meier-Schellersheim

**Affiliations:** Computational Biology Section, Laboratory of Immune System Biology, National Institute of Allergy and Infectious Diseases, National Institutes of Health, Bethesda, MD 20892; Department of Microbiology and Immunology, University of Maryland School of Medicine, Baltimore, MD 21201; Cellular Biology Section, Laboratory of Viral Diseases, National Institute of Allergy and Infectious Diseases, National Institutes of Health, Bethesda, MD 20892; Virginia G. Piper Center for Personalized Diagnostics, The Biodesign Institute, Arizona State University, Tempe, AZ 85287; Center for Vaccine Development and Global Health, University of Maryland School of Medicine, Baltimore, MD 21201

## Abstract

Declining sequencing costs coupled with the increasing availability of easy-to-use kits for the isolation of DNA and RNA transcripts from single cells have driven a rapid proliferation of studies centered around genomic and transcriptomic data. Simultaneously, a wealth of new techniques have been developed that utilize single cell technologies to interrogate a broad range of cell-biological processes. One recently developed technique, transposase-accessible chromatin with sequencing (ATAC) with select antigen profiling by sequencing (ASAPseq), provides a combination of chromatin accessibility assessments with measurements of cell-surface marker expression levels. While software exists for the characterization of these datasets, there currently exists no tool explicitly designed to reformat ASAP surface marker FASTQ data into a count matrix which can then be used for these downstream analyses. To address this, we created CountASAP, an easy-to-use Python package purposefully designed to transform FASTQ files from ASAP experiments into count matrices compatible with commonly-used downstream bioinformatic analysis packages. CountASAP takes advantage of the independence of the relevant data structures to perform fully parallelized matches of each sequenced read to user-supplied input ASAP oligos and unique cell-identifier sequences.

## INTRODUCTION

The experimental workflow of ASAPseq (ATAC with Select Antigen Profiling by sequencing) follows the basic steps of a next-generation, single cell sequencing experiment [1, 2] and we touch on them only briefly. First, nucleic acids from individual cells are partitioned out using a range of experimental protocols [3]. These nucleic acids can, for instance, be cytosolic or nuclear RNA transcripts (RNAseq), accessible genomic DNA (ATACseq), or oligos attached to antibodies labeling surface proteins (CITE/ASAPseq). Next, these partitioned nucleic acids are typically amplified, and adapter sequences either for sample/cell identification or for sequencer-compatibility are added. Sequencing then follows, and the resultant data is transformed into sample-specific FASTQ files. After this FASTQ generation step, the downstream analyses can diverge somewhat depending on the assay [4, 5].

In RNAseq, ATACseq, CITEseq, and ASAPseq applications, the next step typically involves the generation of a count matrix through the alignment of reads to a given reference. While straightforward conceptually, this task is computationally expensive for RNAseq and ATAC-seq specifically, where reads must be mapped to entire genomes. Nonetheless, highly efficient tools exist for completing this task, including the widely used CellRanger software [6], and the open-sourced packages Kallisto/Bustools [7,8] and Salmon [9]. These powerful tools can map reads and identify cellular identifiers (cellIDs) from millions of transcripts across thousands of cells in a matter of hours. While these tools are well-tested for the analysis of RNAseq, CITEseq, and ATACseq data, their application to ASAPseq is less well documented.

ASAPseq is a recently developed approach for quantifying cell surface markers while simultaneously assessing chromatin accessibility of single cells [1, 2]. From a conceptual standpoint, the ASAPseq workflow is directly analogous to that of CITEseq [10]. Antibodies specific to a defined set of surface markers are labelled with unique oligos, which are selectively amplified and sequenced separately from the extracted cellular nucleotides of interest. The downstream analysis of these data should be much simpler than RNAseq and ATACseq; rather than aligning reads to a genomic reference, reads are aligned to a discrete list on the order of 100 sequences to generate the final count matrix. However, a current limitation of CellRanger, the most commonly used processing software, is that it does not support ASAPseq analysis. While Kallisto and Salmon have been reported to be capable of generating ASAPseq count matrices [7–9], at present there is no documentation written specifically for this application. Further, the requirement of a sufficient computational background to compile C/C++ code may exclude a wide range of researchers from using these methods.

To address these issues, we created CountASAP, an easy to install, Python-based tool with support for generating a count matrix from ASAPseq FASTQ files (Figure 1). Here, we provide benchmarking of CountASAP using both ASAPseq and CITEseq datasets. As mentioned previously, ASAPseq and CITEseq datasets are fundamentally similar, allowing us to demonstrate the feasibility of using CountASAP on ASAPseq data while utilizing CITEseq data for a more robust benchmarking against existing methods. We find that CountASAP is capable of properly converting ASAPseq FASTQ files into count matrices, with biologically consistent results between surface markers and chromatin accessibility readouts from ATAC-seq data. Further, CountASAP compares well with CellRanger, the current gold standard in the field, in the analysis of CITEseq data. CountASAP shows exceptional correlation with CellRanger counts on a per-marker per-cell level, and can run efficiently on a standard laptop. With its ease of installation and user-friendly documentation, CountASAP can be quickly incorporated into existing single cell profiling workflows moving forward.

**Figure 1.**
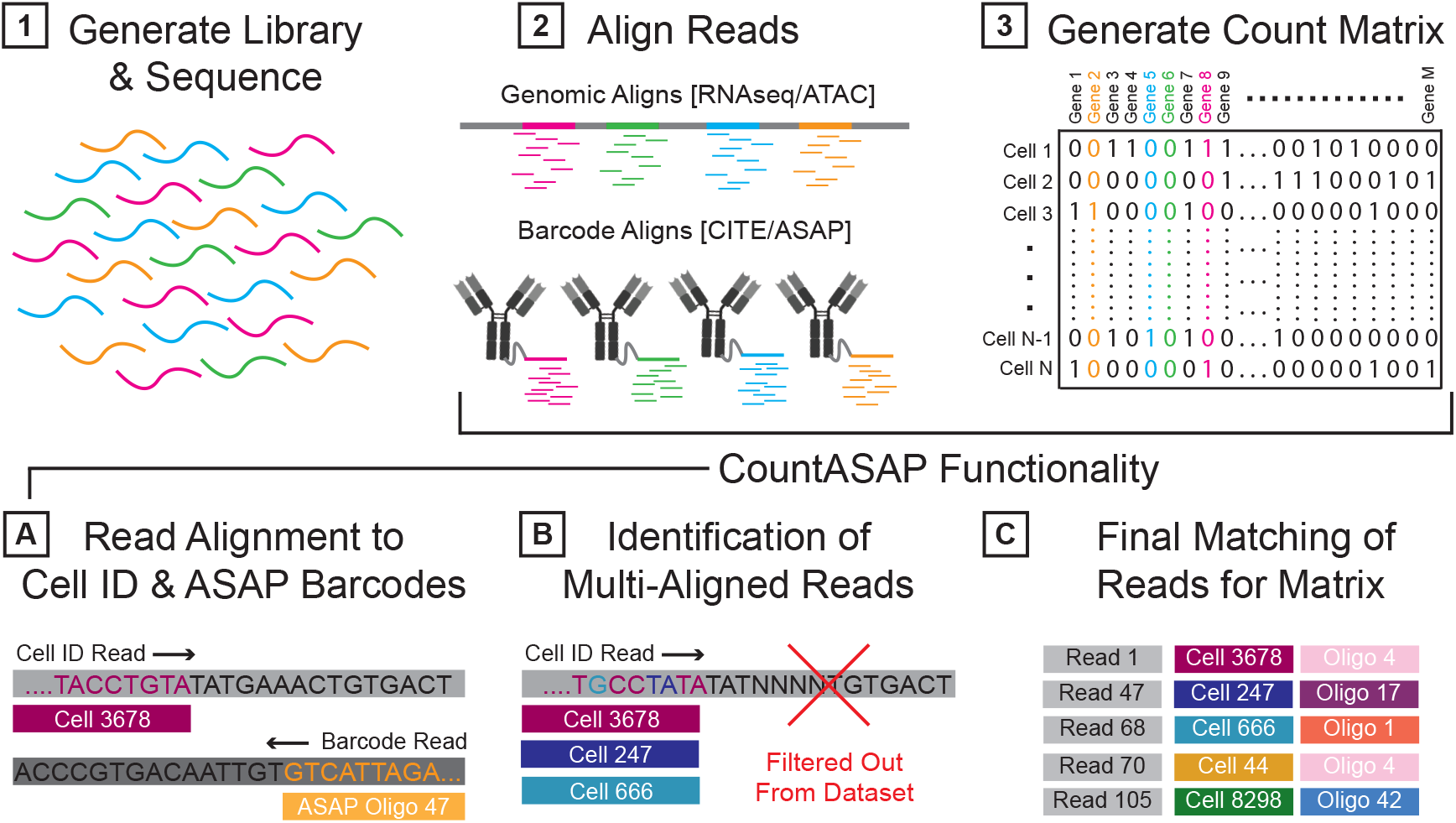
Transforming single cell sequencing experiments into datasets compatible with commonly used analysis pipelines follows a consistent workflow. Libraries are first generated from a given experimental protocol (i.e. RNA/ATAC/CITE/ASAPseq) and sequenced (step 1). Raw reads from sequencers are often output as .BCL files and converted to .FASTQ files. Ultimately, reads from these .FASTQ files must be aligned to some reference (step 2) to generate a count matrix (step 3). In ASAPseq applications, CountASAP can be used to accomplish this task. First, the reads containing cell identification information and ASAP oligo information are isolated and individually aligned to provided lists containing all possible cell identifier and ASAP oligo sequences (step A). CountASAP then identifies those reads that are aligned to multiple references (such as multiple cells or multiple ASAP oligos) and filters these out from the dataset (step B). Final matches of each read with their respective cell identification and oligo identity are compiled in a list, and this list is used to generate the final count matrix (step C).

## RESULTS

To test the capabilities of the CountASAP software, we generated two independent single cell datasets isolated from mice. Starting with 20,000 cells isolated from mouse spleen, we created single cell suspensions which were subsequently profiled using either CITEseq or ASAPseq protocols (See Methods). Both datasets were first processed with CellRanger [6] to create a baseline for comparisons and to identify cellIDs (See Methods), as CountASAP is importantly only for processing surface expression data, not genomic or transcriptomic data. Due to the rarity of surface-marker profiling experiments without corresponding genomic or transcriptomic data, this represents the most-likely use case for CountASAP. All benchmarking comparisons between CountASAP and CellRanger were performed using CITEseq data, due to the lack of ASAPseq support in CellRanger.

### Benchmarking CountASAP Performance

CountASAP was created with the intention of generating a lightweight program compatible across operating systems and with a range of available computing power. The test surface marker CITEseq dataset was collected across four lanes, with two reads in each lane, for a total of 6GB of test data. A total of 24,638 unique cell identifiers (cellIDs) and 211 surface markers provided the maximal dimensions of the final output matrix. With these parameters in mind, the same data was processed using both CountASAP and CellRanger, with quantifiable metrics shown in Table 1.

**Table 1:**
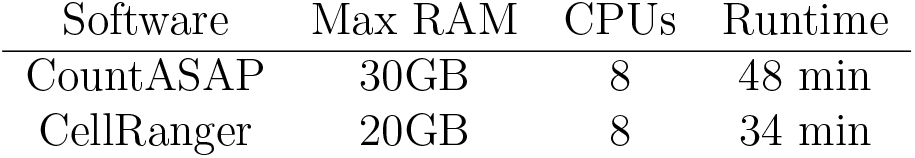
A quantification of the relative memory usage, available CPUs, and average runtime of both CountASAP and CellRanger processing of the same datasets.

Importantly, CountASAP was tested on both a standard laptop (8 cores, 32GB RAM, MacOS) and a Linux machine (32 cores, 256GB RAM), while CellRanger was run on a high-performance compute cluster due to incompatibility with MacOS. We see that overall, memory requirements are only somewhat higher in CountASAP, with some dependence on processing parameter selections (See Methods). CellRanger likewise displays an advantage in the overall time required to fully process the test dataset. We note, however, that the modest disadvantages CountASAP has in memory requirements and speed are a necessary trade-off in making the software user-friendly, easy to install, and compatible across operating systems. Importantly, the final outputs are identical from both programs, in the form of the aforementioned sparse matrix with dimensions determined by the number of cellular identifiers and the number of surface markers in the experiment.

### Identifying Proper Defaults

Having confirmed that the resource requirements and runtime of CountASAP were comparable to CellRanger, we next aimed to identify proper defaults for the software. Given that next-generation sequencing experiments can generate data with a wide range of quality, sequence-processing software needs to be able to handle potential sequencing errors. In ASAPseq (and CITEseq) experiments, the three key components necessary to generate the final count matrix are the cellID, the unique molecular identifier (UMIs), and the oligos conjugated to surface-marker binding antibodies. The unique sequences in these three components are generated such that they are less susceptible to misidentification due to sequencing errors [11], but software that does not allow for base-pair mismatches between the reference sequence and the experimentally identified sequences may severely under-count the identified surface markers associated with each cell, reducing the resolution of the experiment. To quantify the role of such base-pair mismatches between reference and experimental data, we again utilized CITEseq data, specifically focusing on the role of base-pair mismatches in identifying duplicated UMIs (Figure 2).

**Figure 2.**
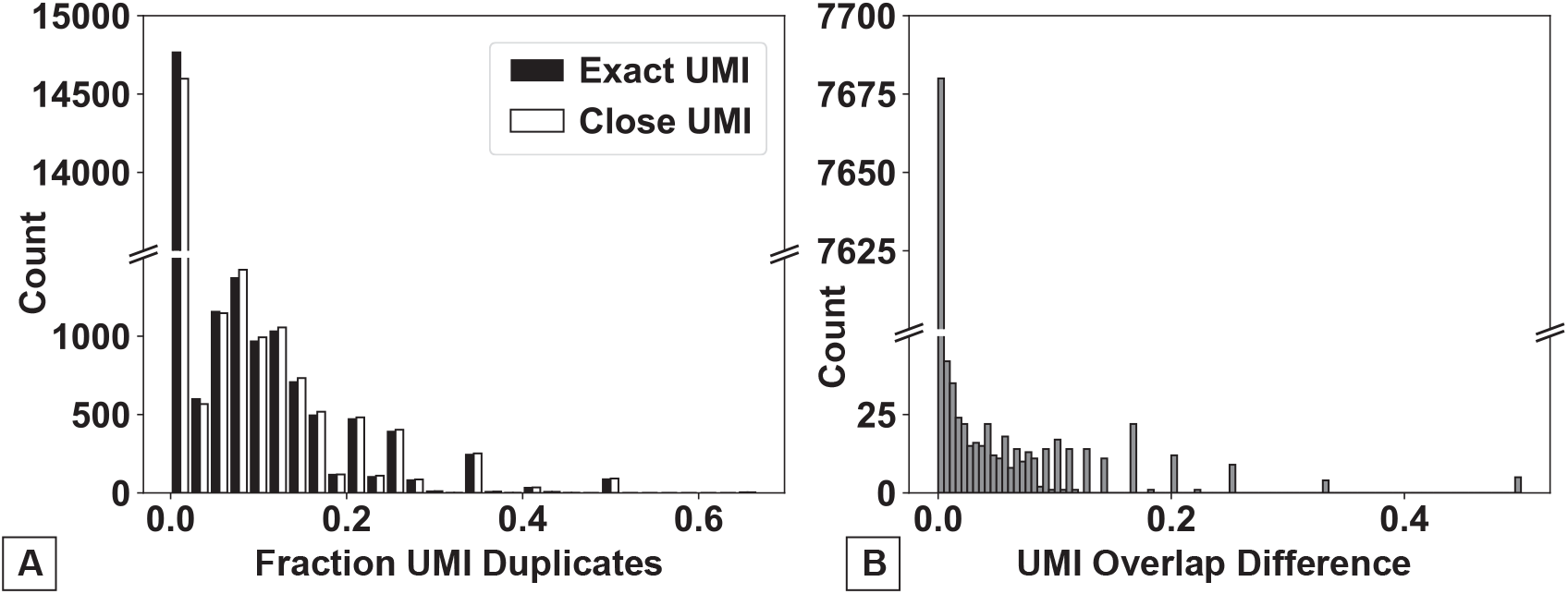
A quantification of the extent of UMI duplication in our CITEseq test dataset and the role of sequencing error on UMI mapping. (A) Duplicate UMIs are identified and quantified using either an exact UMI match (black bars, Exact UMI) or a 1bp mismatch (white bar, Close UMI). Quantification is provided as a histogram of the fraction of the total counts for a given UMI.(B) A histogram of the normalized difference between the “Exact” and the “Close” methods of UMI identification, after the removal of UMIs that have 0 identified duplicates using both methods. Normalization is a simple weighting of the difference in duplicate identification by the total number of total counts of a given UMI. In both figures, note broken y-axes, highlighting order of magnitude difference between zero and non-zero duplicate counts.

We find that in our test CITEseq dataset, the majority of UMIs are not duplicated (approximately 75%, Figure 2A). Of those that are duplicated, only 20% or less of the total UMI counts are duplicates in the majority of instances. Further, we find that duplicate removal using either exact UMI matches or 1bp mismatches does not significantly alter the total number of duplicate UMIs removed. For over 95% of UMIs, using either exact UMI or a 1bp mismatch as the cutoff for identifying duplicates does not result in any change in the number of identified duplicates (Figure 2B). As such, we set the default CountASAP for UMI duplication removal to only drop exact matches from the dataset. Importantly, functionality is maintained for more stringent UMI removal per a user defined basepair mismatch.

The remaining software defaults, specifically the number of basepair mismatches allowed to bin a given sequence with a given barcode or cellID, are set based on the standards in the field. Previous research has suggested that allowing for basepair mismatches over 1 leads to marginal transcript recovery [8]. As such, the standard tolerance for basepair mismatch is set to 1 for non UMI duplication identification. Similar to the UMI duplication function however, users can toggle the mismatch tolerance at the start of each CountASAP instance, if desired.

### A Comparison of CountASAP Outputs to Existing Software

After benchmarking CountASAP and the identification of proper defaults for individual runs, the final required quality control is a direct comparison of the software outputs. After assigning individual reads to a given cellID and antibody oligo barcodes, removal of duplicate UMIs, and dropping of double-counted reads, a final count matrix is generated. The shape and relative sparsity of the matrix is dependent upon the experimental design, but in our test CITEseq dataset 13.8% of the final matrix was nonzero, for a total of approximately 340,000 entries. Of these, 52% of the nonzero entries had only 1 read. Of the remaining 48% of nonzero data with more than 1 read, 99% of the data had under 100 reads. We directly compare these counts identified by CountASAP to the final CITEseq count matrix generated by CellRanger (Figure 3).

**Figure 3.**
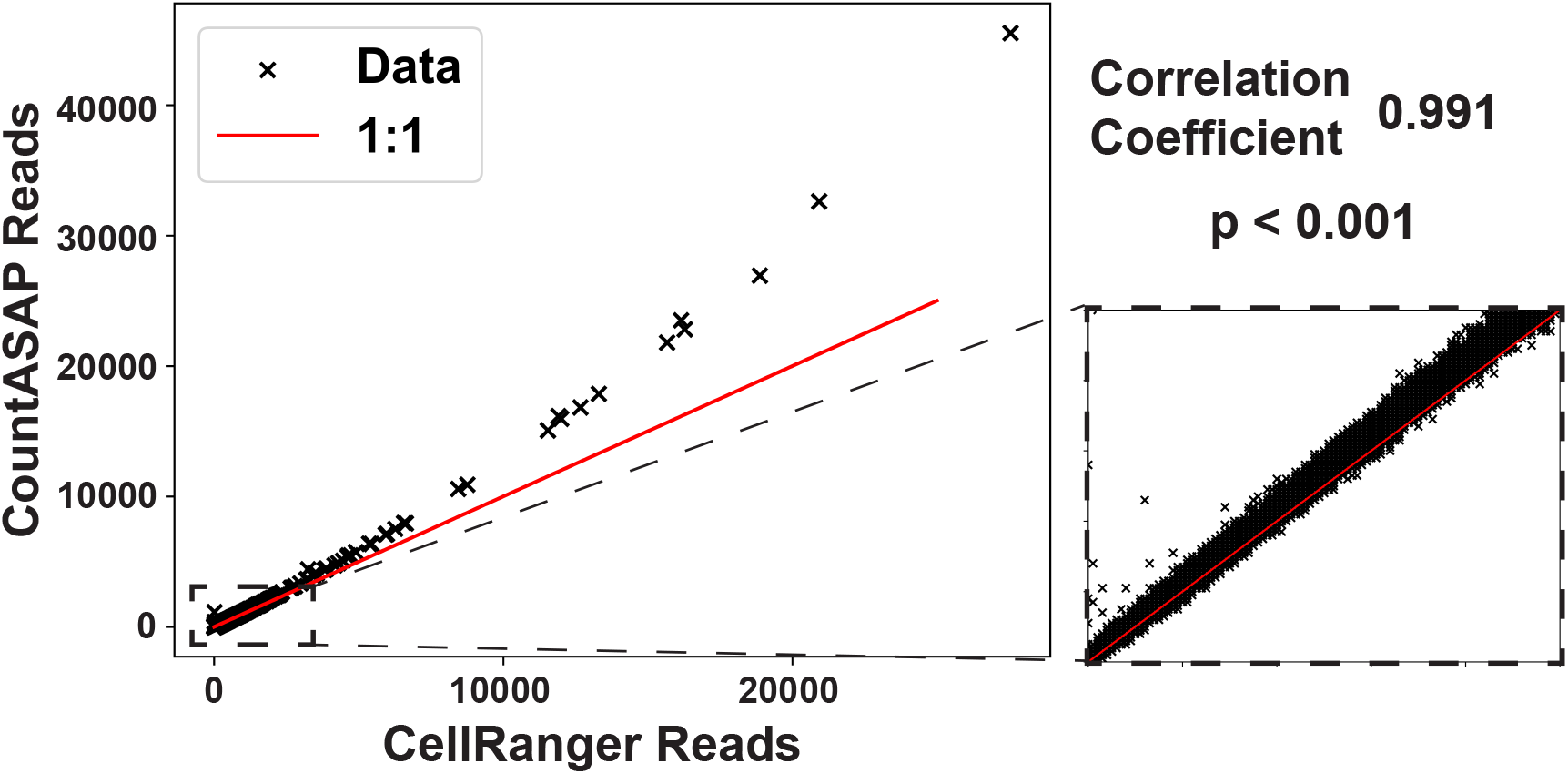
A direct comparison of final matrix outputs generated from CITEseq data by CountASAP and CellRanger. Each data point represents the raw counts for a single antibody oligo and a single cellular identifier. As such, each individual identified cell is represented by 211 points in this plot, corresponding to the total number of CITEseq oligos in the reference list. Inset shows a zoom-in of the region of 0 to 100 CountASAP and CellRanger reads, which includes 99% of nonzero data points. Reported correlation coefficient is for all nonzero data points. Red trend line gives a 1:1 correlation between CountASAP and CellRanger read counts.

We see that overall, CountASAP counts correlate exceptionally well with CellRanger counts. Large outliers from the 1:1 trend line are likewise large outliers in the number of total reads. The zoomed in inset is representative of the vast majority of the entries in the final count matrix, with countASAP identifying roughly 10%-20% more reads per cellID-CITEoligo pair than CellRanger. As the number of reads increase, there is a slight drift towards more total identified reads in CountASAP compared to CellRanger. The strong correlation between the two final count matrices is suggestive of the ability of CountASAP to capture the important biological features of a given CITEseq experiment, and potentially shows that CountASAP can assign reads more efficiently than the current standard in the field.

### Generating Count Matrices of ASAPseq Data

Due to the dearth of software explicitly designed for processing of ASAPseq data, comparisons thus far have relied entirely on CITEseq data. As a final proof of principal of the proper function of CountASAP to reliably map reads from antibody-conjugated oligos to cellular identifiers associated with ATACseq data, we compare the consistency of reads from biologically distinguishing markers in both the chromatin accessibility data and surface marker data. We chose standard markers with the strongest expression and clearest biological function, namely CD4, CD38, and CD11 as surface markers and CD4, Pax5, and S100A9 as genomic markers to identify T cells, B cells, and myeloid cells, respectively.

We first process the ATACseq data using Signac [12], and project this high-dimensional chromatin accessibility data into a two-dimensional space using uniform manifold approximation and projection (UMAP) [13], without the additional processing of a principal component analysis (PCA) step. Removing this often-used PCA step projects the single cells in our ATAC dataset into large clusters with broad biological similarity (Figure 4A). Based on a survey of common markers, we assign a broad classification to each cluster identified by a K-means algorithm (N_*Cluster*_ = 4), identifying two T cell clusters, one B cell cluster, and one myeloid cell cluster. Importantly, the projection steps and clustering are both done without including the surface marker data. In the final comparison step, we see that the normalized expression levels of each marker, chromatin or surface, are highest in the expected cellular clusters (Figure 4B). We find that overall, CountASAP is able to properly assign ASAPseq reads to match the underlying biological context provided by the epigenetic data.

**Figure 4.**
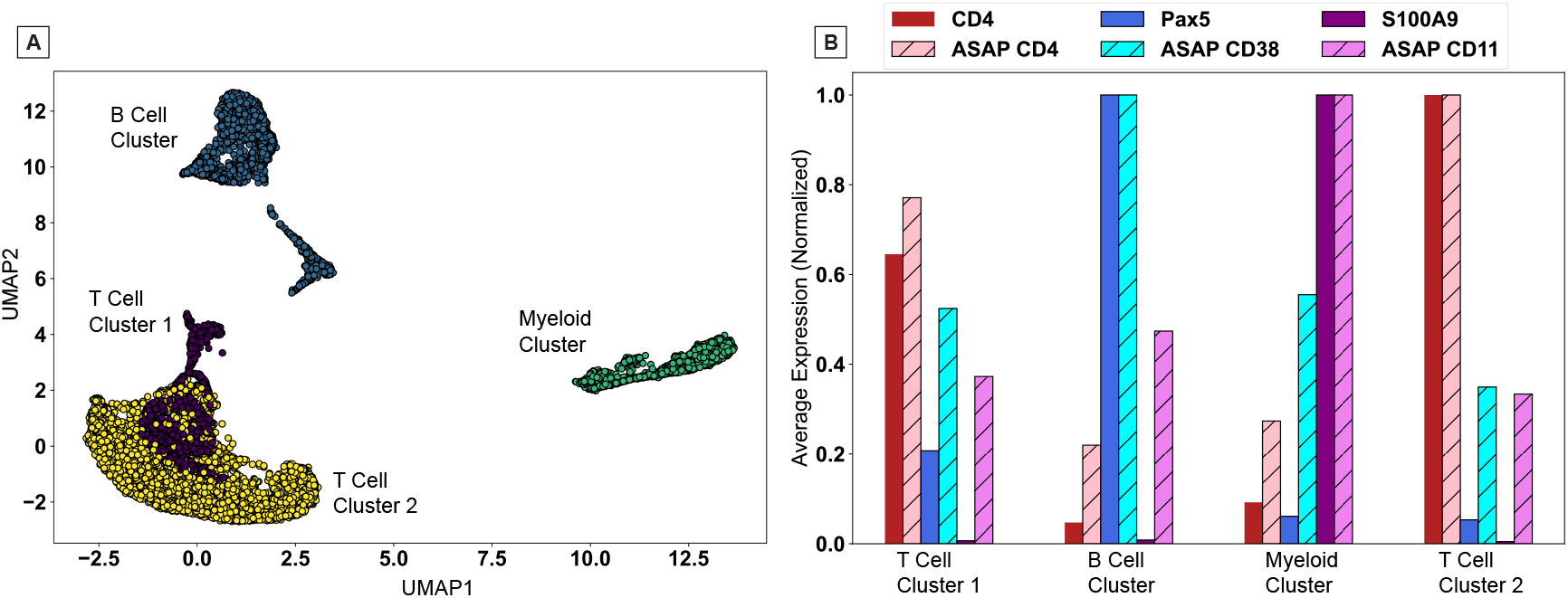
A test of the biological consistency of CountASAP-identified surface marker counts with ATACseq data. (A) UMAP projection of single-cell chromatin accessibility data generates large clusters of broadly biologically similar cells. Colors represent clusters identified using a K-means algorithm, with the number of clusters set to four. (B) Average expression of either chromatin accessibility counts (solid bars) or surface marker counts (striped bars) for each cluster identified in (A). Counts are averaged over entire clusters, and then normalized to the maximum average count across the four clusters.

## DISCUSSION

Alignment software have historically been written in C/C++ to maximize processing speed and parallelize the analysis where applicable. However, code compilation and the required handling of potential incompatibilities that may arise during compilation can create nontrivial obstacles. This issue is strongly alleviated in Python, which is distributed in packages through large repositories such as Conda and PyPi. Additionally, the aforementioned speed advantages accompanying most C/C++ programming are somewhat mitigated by Cython, which some Python packages utilize to rapidly translate and execute code in C [14]. CountASAP takes advantage of such packages, utilizing the easy to execute nature of Python code and the speed of C to create a more user-friendly software for the conversion of ASAPseq FASTQ data into a Seurat- or Signac-readable count matrix.

CountASAP is explicitly designed for the analysis of ASAPseq data, and the only software with a thorough documentation guiding users with step-by-step instructions for this critical pre-processing step. Importantly, this ease of use comes with a minor reduction in performance. While CellRanger, the most commonly used preprocessing software for single cell data processing, is faster than CountASAP, the improvements are marginal for surface marker data. Importantly, CountASAP can be run on a standard laptop running Mac or Windows OS, while CellRanger requires a high-performance computing cluster or Linux machine. In addition to the favorable benchmarking to existing software, the final count matrices generated by CountASAP correlates exceptionally well with those generated by CellRanger, suggesting that no biological information is lost when moving to this open-sourced solution for mapping cell surface marker reads in single cell experiments.

While the majority of the test data presented here are from CITEseq experiments, we were able to show that CountASAP can create count matrices for ASAPseq experiments that show biological consistency between surface markers and chromatin accessibility. We have used CountASAP in the processing of data for our own experimental datasets, and have found that the outputs are compatible with commonly used packages such as Seurat, Signac, ScanPy, and AnnData [15–18].

## METHODS

### Conceptual Overview

Sequencing data in ASAP experiments on the Novaseq S4 platform are typically deposited as four reads, three of which contain information about the ASAP oligo sequenced, the cellular identifier associated with this oligo, and the barcode index. The fourth read contains additional indexing information. CountASAP takes two of these reads as input, identifying the read with oligo identities and the read with cellular identifiers (cellIDs). The user then supplies the ASAP oligos used in the experiment, as well as the expected cellIDs. The list of expected cellIDs can either come from a “whitelist” provided by the manufacturer of the sequencing reagents, or from the cell identification outputs of accompanying ATACseq analysis. ASAP reads that do not match either this whitelist or a list of identified cells are excluded from the final count matrix.

These inputs are then processed and re-formatted into Python lists of strings. From there, the RapidFuzz package is used to quickly match each read to the list of provided ASAP oligos. RapidFuzz provides tunable algorithms for partial matches of strings [19]. In CountASAP, we utilize the RapidFuzz cdist function, which parallelizes and executes the string-matching task in compiled C++. We note that the dataset sizes involved in ASAPseq and CITEseq experiments are too large for RapidFuzz, and must be split into reasonably sized chunks for efficient processing. In this manuscript, we use a chunk size of 100,000 sequences, but this is included as a tunable parameter for users. Cdist further allows for specification of the distance metric used to calculate string similarity. For cellIDs, a Hamming distance [20] is used, while a normalized edit distance ratio is used for the ASAP oligo matching. CountASAP then allows for user-specification of the cutoffs for each of these match functions. The default Hamming distance cutoff is set to 1 mismatch, while the normalized edit distance ratio is set to 0.93, or 1 mismatch with the 16 basepair oligo. These are the recommended cutoffs, as previous research has suggested allowing for greater mismatch leads to marginal transcript recovery [8].

The final step in CountASAP is the removal of duplicated unique molecular identifiers (UMIs). There are two options for the removal of these duplicate UMIs; the faster function removes all exactly duplicated cellID + UMI reads, while the more robust function again utilizes the RapidFuzz cdist function to identify close matches. The faster duplicate UMI removal function is the default in CountASAP. As discussed in the Results, there are marginal differences between exact UMI matches and single basepair mismatches. In the last step, unmatched and duplicated UMI reads are then dropped from the dataset, and the final count matrix is generated from the matched strings. Reports of doubly identified cellIDs or ASAP oligos are included at the end of each instance, so users can identify the errors that may arise from low-quality sequencing data or departures from default mismatch settings. This final count matrix is then formatted identically to CITEseq count matrices currently output by software like CellRanger, and is compatible with commonly used downstream analysis packages.

### Pre-Processing of Data

In this manuscript, both the CITEseq benchmarking dataset (Table 1, Figure 2, Figure 3) as well as the ASAPseq test dataset (Figure 4) required some pre-processing using existing software. Both datasets included complementary transcriptomic or epigenetic data coupled with surface marker expression data, and were first processed with CellRanger to identify cellIDs. CITEseq pre-processing utilized CellRanger version 7.1.0 and the “cellranger multi” instance. ASAPseq pre-processing (ATAC data only) was done with CellRanger-atac version 2.0.0 and the “cellranger-atac count” command. Cellular identifiers were isolated from each of these outputs and used as inputs for CountASAP.

ATACseq data required further processing, especially in the comparison of epigenetic markers with surface markers. We follow the standard processing recommended by Signac [12] to normalize raw counts output by CellRanger. We use standard quality control filters on the cells kept in the datasets, requiring cells have between 3,000 and 30,000 reads, more than 15% of reads within peaks, a blacklist ratio less than 0.05, a nucleosome signal less than 4 and a TSS enrichment greater than 2. Regions of the mouse chromosomes are annotated using EnsDb.Mmusculus.v79, and the peaks are saved and exported in matrix form. From there, data are loaded into a custom Python script for downstream analysis to create figure four. All pre-processing and figure-generation scripts are included in the CountASAP GitHub.

### Experimental Data Acquisition

Experimental protocols for CITEseq and ASAPseq data acquisition followed a similar design, and are outlined in parallel here. In both experiments, dissociated single cell suspensions from each mouse were dispensed into two sets of 96-well plates, with each well containing approximately 150,000 cells. Both sets were blocked with an anti-mouse CD16/32 antibody (Trustain FcX Plus: BioLegend) to prevent non-specific binding via the Fc receptor. After blocking, one set was stained with Total Seq-B anti-mouse hashtag antibody (BioLegend) for ASAP, and the other was stained with TotalSeq-C anti-mouse hashtag antibody (BioLegend) for CITEseq. Hash-tagged cells were then pooled together (approx. 2 million cells) and stained with a mixture of sixty-two TotalSeq-B anti-mouse surface protein panel antibodies (for ASAP) or seventy-hour TotalSeq-C anti-mouse surface protein panel (for CITEseq), specific to certain cell lineages, at concentrations ranging from 12.5ng to 100ng per pooled sample. Stained cells were washed with Cell Staining Buffer (BioLegend). Cell numbers were counted, along with viability.

For CITEseq preparation, we targeted a maximum input of 20,000 cells using “Chromium Next GEM Single Cell 5’ HT Reagent Kits, v2” (10x Genomics) and followed the manufacturer’s protocol (CG000424 Rev C) without any modification. For nuclei preparation (ASAP), we followed the protocol from the NYGC Innovation Lab [1] with a slight modification. Surface protein stained cells were fixed in a 1% paraformaldehyde solution in phosphate buffered saline (PBS) for 10 min at room temperature and washed with cold PBS. These fixed cells were resuspended in LLL Lysis buffer (10mM Tris-HCl (pH 7.5), 10mM Sodium Chloride (NaCl), 3mM Magnesium chloride (MgCl_2_), 0.1% NP40, 1% bovine serum albumin (BSA)) and incubated for 3 min on ice, washed with Wash buffer (10mM Tris-HCl (pH 7.5), 10mM NaCl, 3mM MgCl_2_, 1% BSA), and finally resuspended in 1x nuclei buffer (10x Genomics). The number of nuclei was counted and loaded onto Chromium Next GEM Single Cell ATAC Reagent Kits v2 (10x Genomics). We followed the manufacturer’s protocol for ATAC library construction. For ASAP, we followed the protocol from NYGC Innovation Lab by adding a bridge oligo B (BOB) at Step 2.1a and saving the supernatant at Step 3.2d in the 10x Genomics’ ATAC protocol (CG000496 Rev B). The saved supernatant was cleaned by using SPRIselect Reagent (Beckman Coulter) with a magnetic separator and proceeded to the index polymerase chain reaction (PCR) with D7xx s primers. The final products (approx. 200 bp) were cleaned up with 1.6x SPRIselect reagent. Both ATAC and ASAP libraries ((or CITEseq surface and transcriptomic libraries) were pooled together and sequenced in Illumina Novaseq 6000.

### Availability and Implementation

CountASAP is freely available on GitHub [https://github.com/ctboughter/countASAP] complete with documentation and sub-sampled test data. For easy installation, the complete Python package is also included on PyPI [pip install countASAP], with instructions within the documentation outlining the steps for creating Python environments.

## Acknowledgements

This work was supported by the intramural program of the National Institute of Allergy and Infectious Diseases (NIAID), NIH. This study was funded by the Defense Advanced Research Projects Agency (DARPA), Assessing Immune Memory (AIM) program under grant HR001121S0037-AIM-FP-009.

